# Optimal forgetting: Semantic compression of episodic memories

**DOI:** 10.1101/2020.05.06.080838

**Authors:** David G. Nagy, Balázs Török, Gergő Orbán

## Abstract

It has extensively been documented that human memory exhibits a wide range of systematic distortions, which have been associated with resource constraints. Resource constraints on memory can be formalised in the normative framework of lossy compression, however traditional lossy compression algorithms result in qualitatively different distortions to those found in experiments with humans. We argue that the form of distortions is characteristic of relying on a generative model adapted to the environment for compression. We show that this semantic compression framework can provide a unifying explanation of a wide variety of memory phenomena. We harness recent advances in learning deep generative models, that yield powerful tools to approximate generative models of complex data. We use three datasets, chess games, natural text, and hand-drawn sketches, to demonstrate the effects of semantic compression on memory performance. Our model accounts for memory distortions related to domain expertise, gist-based distortions, contextual effects, and delayed recall.

**Author summary:** Human memory performs surprisingly poorly in many everyday tasks, which have been richly documented in laboratory experiments. While constraints on memory resources necessarily imply a loss of information, it is possible to do well or badly *in relation* to available memory resources. In this paper we recruit information theory, which establishes how to optimally lose information based on prior and complete knowledge of environmental statistics. For this, we address two challenges. 1, The environmental statistics is not known for the brain, rather these have to be learned over time from limited observations. 2, Information theory does not specify how different distortions of original experiences should be penalised. In this paper we tackle these challenges by assuming that a latent variable generative model of the environment is maintained in semantic memory. We show that compression of experiences through a generative model gives rise to systematic distortions that qualitatively correspond to a diverse range of observations in the experimental literature.

## Introduction

It has long been known that human memory is far from an exact reinstatement of past sensory experience. In fact, memory has been found surprisingly poor for even very frequently encountered objects such as coins [1], traffic signs [2] or brand logos [3]. Rather than being random noise however, the distortions in recalled experience show robust and structured biases. A great number of experiments have shed light on systematic ways in which the distortions in recalled memories can be influenced both by past and future information, as well as the context of encoding and recall. Canonical examples of past knowledge influencing recall include experiments of Bartlett [4] where for folk tales recalled by subjects of non-matching cultural background, the recalled versions were found to be modified in ways that made the stories more consistent with the subjects’ cultural background, leading to the suggestion that memory seems to be more reconstructive than reproductive. Various manipulations of the encoding context have been shown to influence the recalled memory; for example, presenting a label or theme before ambiguous sketch or text stimuli modulates both recall accuracy and the kinds of distortions that appear in the recalled memory [5–7]. Finally, paradigmatic examples of memory disruptions due to information obtained after the experience being recalled include post-event misinformation [8], imagination inflation [9], hindsight bias [10], or leading questions [11]. The rich set of systematic distortions provides insights into the principles governing memory formation, which ultimately provides a means to predict how experiences are transformed in memory.

Traditionally, systematic biases in recalled memories have been interpreted as failures, confabulations of an unreliable memory system [12]. In contrast, adaptive accounts have been proposed that view these biases as being regrettable but necessary byproducts of adaptive processes in the brain such as generalisation, fast recognition or creativity [13–15]. Going even further, Bayesian accounts of reconstructive memory argued that in some cases, the memory distortions can be adaptive even if the goal is accurate recall, since previous knowledge can be used to correct for inaccuracies and fill in missing details in incomplete memories [16–18]. The Bayesian account provides a principled way of decoding memory traces by combining prior statistical knowledge with noise-corrupted information retained from the observation. However, it leaves a fundamental question open: in a normative model of memory what information needs to be retained and what pieces of information should be sacrificed to satisfy constraints on memory resources?

In this paper we argue that viewing the transformation of sensory experiences into memory traces as compression provides a normative framework for memory distortions. Specifically, we consider lossy compression, a form of compression where limited-capacity encoding is achieved at the price of imperfect decodability of original data, to characterise information loss during encoding. We point out that by adapting the mathematical framework for lossy compression, called rate distortion theory, to the constraints faced by the brain, we obtain semantic compression. Key to semantic compression is the assumption that a generative model of the environment is maintained in the brain. This generative model describes how the observed statistics of the environment has been generated from variables not directly observed through our senses [19]. According to semantic compression, it is the latent variables of this generative model of the environment that are used to compress experiences. If memory is optimised for natural observation statistics, then assessing the predictions of lossy compression regarding memory distortions requires generative models capable of handling such complex structured data. Constructing such generative models and performing inference in them can be challenging therefore we capitalise on recent advances in machine learning: we use variational autoencoders to learn approximate generative models of structured data [20]. Importantly, a form of variational autoencoders, the beta-variational autoencoders can be viewed as a variational approximation to rate distortion theory, and we use this link between rate distortion theory and generative models to provide a theoretical framework for a unifying explanation of a large body of experimental data in the domain of memory distortions.

In the following, we introduce the theoretical background for semantic compression and then apply semantic compression to three different domains in memory distortions. First, we introduce basic concepts of rate distortion theory. Second, we introduce variational autoencoders and their relationship to lossy compression. Next, a paradigmatic example of memory distortion induced by past experience, domain expertise is investigated. In the coming section we discuss contextual effects through semantic compression. Finally, we discuss how gradual change of compression as time progresses incurs changes in memory traces.

## Theoretical framework

### Rate distortion theory

The branch of information theory that deals with data compression where information is lost during the process is rate distortion theory (RDT). According to RDT, a compact code is constructed that can be used to encode any data point in the dataset. A central insight of RDT is that there is no single optimal encoding: a trade-off emerges between the memory resources (rate, *R*) that are used for storing a given observation, i.e. the length of the code and the expected amount of distortion (*D*) in the recalled memory, i.e. the reconstruction of the original data from the stored code. Any given compression algorithm can be characterised by the trade-off it makes between these two quantities, and thus defines a point in the rate distortion plane (RD plane). An encoding, *Q*, can be improved by decreasing the expected distortion, *D*_*Q*_, without increasing the rate, *R*_*Q*_. Thus, the best encoding under any given memory resource constraint *R* is the one that minimises the expected distortion when the rate is maximised at a specific value, *R*:

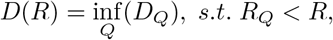

or alternatively, by achieving a lower rate without increasing the expected distortion. Achieving the lowest possible distortion for each possible rate traces out the RD curve, establishing the range of possible optimal encoding schemes for a given distribution over observations.

Under any given encoding, the distortion of an individual observation 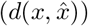 between the observation (*x*) and its reconstruction 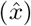 can change across observations. The distortion term measures the expected amount of error that the encoding algorithm makes over the whole set of observations. The function 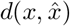 characterises how acceptable the distortion is and thereby defines an ordering across pairs of observations and their reconstructions, which defines the contribution of an observation to the the distortion term of RDT, 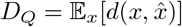. An optimal lossy compression algorithm will selectively prioritise information such that alterations that are inconsequential according to this measure are discarded first.

An alternative formulation of finding the best distortion, *D*_*Q*_, under the constraint of limited rate, *R*_*Q*_ can provide additional insights into the continuum of solutions obtained at different rates. The Lagrange-multiplier formalism is used for constrained optimisation of the distortion such that instead of minimising *D*_*Q*_ one needs to minimise

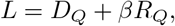

where the constraint of fixed rate when optimising the distortion is formulated through the Lagrange-multiplier *β*, which sets the trade-off between two terms. This formulation of the objective is only applicable when the RD curve is strictly convex but this requirement is often fulfilled in practical cases. According to this Lagrangian formulation, any compression method can be associated with a point on the RD plane, with optimal algorithms lying on the curve. Every point on it can be identified with a single value of *β*, which is the local slope of the curve. Thus, *β* directly corresponds to a particular point on the rate-distortion trade-off continuum: for example a high value of *β* is associated with strong compression, yielding a low rate but high distortion.

While RDT provides a normative framework for lossy compression, the errors or compression artefacts resulting from traditional lossy compression algorithms of images or video (such as block boundary artefacts characteristic of JPEG) look qualitatively different from the errors committed by human memory [21, 22]. If one intends to use RDT as a normative framework of human memory errors then such mismatch ostensibly casts doubt on the applicability of the framework to memory phenomena. However, compression of sensory experience for the human brain is characterised by a number of constraints which distinguish it from the traditional problem of compression of image and video data. We propose to accommodate these constraints in the framework of RDT to obtain semantic compression.

### Semantic compression

A fundamental difference between RDT and the form of compression required by the brain is that while the former produces optimal reconstructions given a known source distribution, the distribution of observations is not known for the brain but has to be learned from experience over time. Specifically, natural observation statistics define richly structured and high dimensional distributions, which have to be learned from a comparatively limited set of observations. In machine learning, this challenge is addressed by generative models. In order to cope with severely limited training data, generative models include inductive biases such as restrictions on hypothesis spaces, priors or hyperparameters. These biases influence decisions regarding what features of the data are generalisable to future observations and what features should be deemed random noise. We argue that since the problem of generalising from a small amount of observations to the true underlying distribution is a fundamental challenge in both generative models and compression in memory, similar inductive biases have to be incorporated in both. Therefore, we propose that the normative approach for adapting RDT to the problem of human memory is through compression via generative models.

Probabilistic generative models have been implicated in understanding human and animal behaviour in a multitude of cognitive tasks [23–26]. For example, perception has been previously cast as a process of unconscious inference, which is aimed at inferring the latent state of the environment based on noisy sensory observations [19, 27]. This inference can be accomplished optimally by inverting the generative model which describes the way latent variables give rise to observations. Another domain where generative models have found support is action planning, where the model is used as an environment simulator to predict likely consequences of actions [28]. Following previous research, we assume that a statistical model of the environment is maintained in semantic memory and formalise semantic memory as a probabilistic generative latent variable model of the environment [29–31]. We argue that semantic memory represents the best estimate the brain has of environmental statistics, and therefore assume that it is this approximate model of the environmental statistics that compression is optimised for.

When the RDT framework is applied to the compression problem faced by the brain, a further issue needs to be considered: RDT does not specify how distortion should be measured, leaving it to be defined by the application. The distortion function represents the agent’s judgements on how relevant particular features of the observation are and efficient compression hinges on selectively retaining this information. Thus, the definition of the distortion function raises the question of what parts of experience are relevant for the human brain. We argue that this problem is identical to that encountered in perception, where computations aim at extracting the latent variables underlying the activity of sensory neurons. In semantic compression the distortion function is defined by the generative model maintained by semantic memory, and the relevant features of observations are those that are extracted into the latent variables of this model. Importantly, such a choice for the distortion function and optimising for the complex structure in natural observation statistics can yield qualitatively different errors from those made by traditional compression algorithms, which only exploit simple, low-level regularities in image statistics.

### Variational approach

Generative models and rate distortion theory are separate frameworks developed with largely different goals, however there has been a flurry of recent work pointing out connections between the two [32–34]. A recent development in machine learning is the introduction of variational autoencoders, which can effectively learn a generative model of complex, high dimensional distributions as well as perform approximate inference over the latent variables. Interestingly, recent studies have established a link between a variant of variational autoencoders and rate distortion theory. Here we briefly describe the variational framework in which rate distortion theory and generative models can be jointly discussed.

Learning probabilistic latent variable generative models of natural stimuli such as images, videos and sound has been a major challenge in machine learning and requires approximate methods. Many of these models utilise latent variables, *z*, to factorise the distribution over observations, *x*. The set of latent variables can be thought of as the factors that contribute to the structure of the input data and constitute a representation of the data, often with lower dimensionality. These latent representations often show desirable qualities such as disentangling independent factors of variation. One of the most successful approaches to learning approximate latent variable generative models is a class of models called variational autoencoders (VAE) [20]. VAEs are capable of jointly learning the parameters of the generative model as well as performing inference by approximating the posterior distribution over latent state variables through variational methods. In variational Bayesian inference the true posterior distribution, *p*(*z*|*x*), is approximated by a distribution *q*_*ϕ*_(*z*|*x*) from a simpler distribution family parameterised by *ϕ*. Once such a distribution family is chosen, the goal is to minimise the dissimilarity between the true posterior and the approximate posterior:

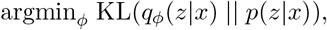

where KL is the Kullback-Leibler divergence, which quantifies the dissimilarity between two probability distributions. While this term cannot be computed directly, it can be shown that maximising the evidence lower bound (ELBO),

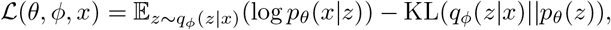

also minimises the KL divergence. In addition to learning to perform accurate inference through optimising the parameters *ϕ*, the generative model is also learned through optimising the ELBO over the parameters *θ*. The generative model consists of the likelihood, *p*_*θ*_(*x*|*z*), describing how the observed variables depend on the latents, and the prior distribution over latents *p*_*θ*_(*z*). The first term of the ELBO is often called the reconstruction term, alluding to the fact that it penalises inaccurate reconstruction of the observation. The second term is usually viewed as a regularisation term, as it penalises deviations from a simple posterior. In VAEs the approximate posterior is typically of a simple form, such as a Gaussian, which is parameterised by its mean and covariance structure. VAEs also utilise amortised inference, where the computation of the approximate posterior is amortised by training a neural network to output the parameters of the approximate posterior on the training set. The output of this *encoder* then produces the parameters of the approximate posterior over *z* in later observations. An extension of VAEs, *β*-VAE, is particularly relevant for establishing a formal connection between generative models and RDT. *β*-VAE introduces a scalar multiplier, *β*, that scales the regularisation term and can effectively trade-off the two terms with the motivation that individual latent variables of the learned latent representation correspond to independent and interpretable sources of variation in the observed data, i.e. encouraging more disentangled representations [35].

To understand the relationship between *β*-VAE and RDT, we turn to a specific formulation of compression called the information bottleneck (IB) method [36]. The IB method extends RDT such that it guides the choice of the distortion function. The IB method introduces the term *relevant information*, the information that we intend to retain after compression. If the goal of compression is to lose information such that estimation of the relevant quantity, *y*, is minimally affected then we can formulate the relevant information as the information in the compressed representation *z* with respect to the relevant quantity *y*, that is the mutual information *I*(*z, y*). Consequently, in the loss function of IB the goal of maximising relevant information is traded off with compressing observations through the latent representation, and the loss to be minimised becomes:

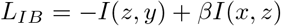

It can be shown that minimising the IB loss function corresponds to a distortion measure that prioritises information that contributes to the prediction of relevant quantity *y* [36]. Although the IB method provides an algorithm for optimising the loss function, it is not feasible to apply to high dimensional naturalistic data. Alemi et al. [32] have shown that the IB objective can be efficiently approximated through variational methods. Importantly, the IB method formally defines a supervised objective since optimisation of the compression is achieved with the objective of optimising for a particular ‘output’ variable *y*, but an unsupervised version of the IB method can be constructed which gives rise to the same objective as the *β*-VAE. In this sense, *β*-VAE can be seen both as a generative model and a lossy compression algorithm: the reconstruction term in the ELBO can be interpreted as the distortion, *D*, and the information limiting regularisation term as the rate, *R*.

The correspondence between RDT and *β*-VAEs highlights the relation of latent variable models to lossy compression. Inferring a posterior over latent variables *z* upon the observation of stimulus *x* amounts to only retaining the statistics of stimuli captured in variations in *z* but discarding those beyond the sufficient statistics of the latent variables.

In summary, we use the framework of VAEs to learn the kind of generative model hypothesised to be maintained by the human brain and we link approximate inference over the latent variables of *β*-VAE to inferences made by humans. We then analyse this model from the point of view of lossy compression, allowing us to model and provide a normative explanation for a large variety of memory experiments.

## Results

### Domain expertise

According to semantic compression, efficient compression hinges upon accurate knowledge of environmental statistics. Since in the case of the brain these statistics are estimated based on experience collected over time, the accuracy of the estimate is expected to increase with the amount of experience within a cognitive domain. As the estimate becomes more accurate, compression becomes closer to optimal and consequently recall errors are expected to decrease. However, this enhancement in recall accuracy is only expected to occur for observations congruent with the statistics of the domain, as a compression algorithm optimised for one distribution will be poorer at encoding observations coming from a different distribution. Assuming semantic compression, constructing artificial stimuli of the same domain but exhibiting statistical structure incongruent with that of earlier experience will increase recall errors.

Chess is an ideal domain for computational analysis of expertise on memory performance due to a number of factors. i) The data is rich, possible configurations are astronomical; ii) chess games trace out a complex subspace of possible configurations; iii) ‘natural’ game statistics is well documented; iv) expertise is graded among individuals, allowing for a more fine-grained analysis of the relationship between expertise and recall performance. We capitalise on these properties of chess to test how expertise relates to memory performance in different conditions.

In a widely studied paradigm in memory research using chess [37], a chess board configuration is presented for less than 10 seconds, after which pieces are removed and subjects are required to reconstruct the observed configuration by placing the pieces on an empty board. Subjects are classified into four skill levels on the basis of their Elo points. Recall performance is measured in two conditions: In the case of ‘game’ (or ‘meaningful’) configurations chess pieces are placed according to states taken from actual games, while in the case of ‘random’ (or ‘meaningless’) configurations positions of chess pieces of game states were randomly shuffled.

We trained a *β*-VAE to learn the distribution of chess pieces during standard chess games downloaded from the FICS games database^1^ (chess-VAE). Briefly, a board configuration was represented as a 64 by 13 element matrix corresponding to the 64 positions and the 13 possible pieces, with an element of the matrix taking one if a particular chess piece appeared on a given position. This input was encoded with the *β*-VAE in a 64-dimensional latent space (for additional details please refer to the Materials and Methods section). In order to capture the varying amounts of experience that subjects have with these statistics, we trained the generative model on varying amounts of chess games, using 0.1% (unskilled) to 90% (most skilled) of the entire training set consisting of approximately 250000 board configurations. In addition to the chess games, in order to mitigate overfitting to a low number of observations for the unskilled model, we augmented the training data with 10000 uniformly random board configurations at each skill level. This data augmentation can be seen as a hand-crafted inductive bias which optimises for a uniform distribution in the low data regime.

Optimising for an uniform input distribution means that the algorithm maintains an ability to reconstruct any possible board configuration equally well, however since overall capacity is limited, this means that no configurations can be reconstructed accurately. In the case of more skilled models the observations overwhelm the prior and consequently the prior has negligible effect. Note that we are taking a conservative approach in training the model, with no explicit instructions regarding the rules of chess or intent to win the game. Explicit knowledge of the rules makes certain configurations impossible or exceedingly unlikely, which can be utilised to aid recall. Nevertheless, reconstructions and unconditional samples show that the model captures an approximate version of these rules. To model the experimental recall setting, we used the inference network of the learned generative model to encode either game or random boards into a latent representation. Then, conditioning on the stored latent state we used the generative model to decode the memory trace into a reconstruction of the chess board configuration (Fig. 1A,B). Reconstructions by the model show the monotonic increase in accuracy for ‘game’ boards as a function of increasing chess skill (Fig. 1C). A similar monotonic increase in recall performance was found in humans (Fig. 1D), where recall performance ranged from around five pieces for amateur players to near perfect reconstruction for grandmasters.

**Fig 1.**
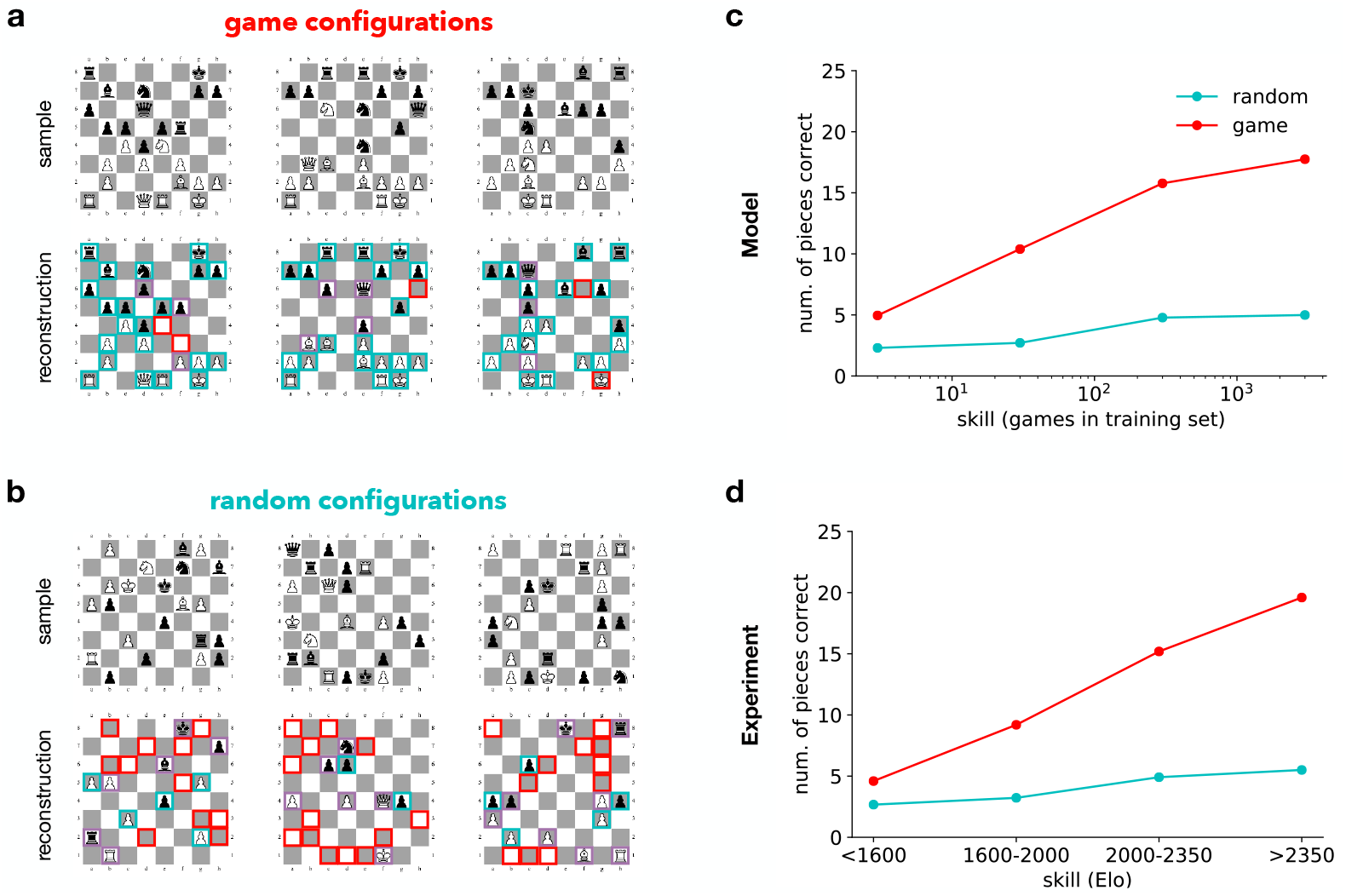
Effect of domain expertise on memorising positions in chess. A: *Top*, Chess board configurations from real game settings (*game configurations*). *Bottom*, reconstructions of the configurations from memory. These configurations are individual samples generated by the expert model based on the encoding of the presented configurations. *Green frames* indicate correctly reconstructed pieces, *red frames* indicate positions where a piece is missing or erroneously appears in the reconstructed game; *purple frames* indicate pieces whose identity is switched in the reconstruction. B: Same as A but instead of game configurations randomly shuffled pieces are presented (*random configurations*) and reconstructed. C: Reconstruction accuracy of the model for game and random configurations as a function of the training size. D: Reconstruction accuracy of human participants as a function of chess skill. Data reproduced from [37].

In the ‘random’ condition, artificial stimuli obtained by randomisation destroys a significant portion of the statistical structure present in the configuration. In the case of ‘random’ boards the chess-VAE displays more errors both in omitted pieces and in exchanged piece identities (Fig. 1B). As a function of expertise, a monotonic increase in accuracy can be observed but with a distinctly smaller slope than in ‘game’ positions (Fig. 1C). In contrast to the ‘game’ stimuli case, for these artificial stimuli the accuracy advantage of skilled human players also shrinks substantially (Fig 1D), meaning that the accuracy advantage of skilled players originates in the statistical structure of the stimuli.

Naively, one might expect that it would be easier to recall boards with only a few pieces, with the underlying assumption that storing the location and identity of each additional piece requires additional memory resources. However, board configurations containing most of the pieces are usually from early in the game, with the configuration strongly constrained by the initial state, whereas boards containing only a few pieces are typically from late in the game less constrained by the starting game setting. Intuitively, differences in compressibility can be understood to arise from the relatively short description length of an early game configuration where one only needs to define the movements of a few pieces relative to the initial state. In summary, increased expertise in the statistics of a particular stimulus set specifically contributes to the enhancement of recall performance, which can be explained by recruiting knowledge stored in semantic memory for efficient compression of data.

### Gist-based distortions

Semantic compression assumes that the statistics of stimuli is learned through a generative model and the latent variables of this generative model determine what features of experience are retained in lossy compression. Ideally, the latent representation that a generative model learns captures factors that explain a large amount of variance in earlier observations, are strongly predictive of future observations and rewards or allow for efficient manipulation of the environment. These latent variables are hypothesised to include lower level acoustic or visual features such as phonemes, or objects as well as abstract concepts such as what constitutes a good chess move or melody. These more abstract latent variables provide a high level, ‘gist’-like description of the experience. By conditioning the generative model on the latent representation, observations that are consistent with the high level description can be generated. While precise details of the episodes will be lost during the encoding and decoding process, lost information can be supplemented by the generative model during decoding. More specifically, the generative model can be used to generate likely values of features for which the observed value was discarded during compression.

One consequence of reconstruction through a generative model is that memory will be sensitive to changes in the observations that affect the latent variables but allow for distortions that do not. At sufficient levels of compression this will result in falsely recognising or recalling items that were not themselves presented, but are conceptually related to items that were.

A second consequence of compression using latent variables of a generative model is that factors that influence the interpretation of the observation, that is the inference of values of latent variables, will also be reflected in the reconstruction. Specifically, in the case of ambiguous stimuli, contextual information influences the inferred latent representation, and consequently distorts the compressed memory by shaping lower-level details in ways that better conform to the shifted latent representation.

We demonstrate these effects in two experimental domains, the delayed recall of lists of words and recall of hand drawn sketches of objects. Note, that it is currently a challenge in machine learning to identify the computational principles that give rise to generative models that decompose observations into latent variables resembling the representations in human semantic memory. A particular advantage of using *β*-VAEs is that one of the main ingredients to achieve learning such a representation is thought to be the principle of encouraging disentangled features. *β*-VAEs have been shown to be able to discover disentangled latent representations from complex data in diverse domains and are therefore a good candidate for investigating the forms of memory distortions that result from the manipulations of latent representations in a domain-general way.

#### Intrusion of semantically related items during recall

One of the most extensively studied paradigms for reliably inducing strong false memories is the Deese–Roediger–McDermott paradigm (DRM) paradigm, where subjects have to recall lists of words. Language is a rich and computationally difficult domain, however recent successes in generative modelling of language suggest that the available size of corpuses makes it amenable to learning in an unsupervised way [38]. An inherent advantage of the DRM experimental setting is that since the recall order of word lists is not constrained, simpler, so called ‘bag of words’ models can be used which are significantly easier to train than text models that can also capture sequential dependencies between words.

In the DRM paradigm [39], subjects are presented with a list of semantically related words. The lists are created by collecting first associates of a particular common word, the lure word, from human subjects. In the experiment word lists are created from associates and presented to participants but the lure word is never shown during the memorisation phase. After a given delay that ranges from minutes to days, subjects have to either recall or recognise the studied words.

In order to learn an approximate generative model for language, we have used Wikipedia excerpts subsampled to 40 words for training a simple VAE architecture (text-VAE). The architecture was similar to the chess-VAE except that the noise model and input representation was adapted to text observations. This architecture has also been analysed previously in the machine learning literature as the Neural Variational Document Model (NVDM) [40]. The model takes text snippets as input to the *β*-VAE in bag of words representation, i.e. each occurrence of a word in a document is counted in a vector of dimension equal to the size of the entire vocabulary. This representation of the input disregards the sequential structure of text. The encoder mapping a document to a latent representation **z** of 100 dimensions consists of two dense layers of 2000 hidden units. The generative model is similarly structured and generates words independently (for additional details, see the Materials and Methods section). Wikipedia was chosen as a large and reasonably comprehensive corpus. After training the text-VAE model, for any presented list of words, a posterior can be inferred, which corresponds to the latent variables that might underlie the observed word list. Nearest associates of the lure words show that the learned representation captures similar statistical relationship between words as the methods used in the original experiments to generate associate word lists (25% of corresponding DRM list words appear in 50 closest associates to lure word in model, for examples see Methods). However, since the statistics of words in an encyclopedia is different from that encountered by a human during his lifetime, the model’s interpretation of certain words can be biased (e.g ‘chair’ is strongly related to ‘organization’ in Wikipedia dataset but not in the DRM word lists).

Generative models of text such as topic models often make the assumption that natural text is concentrated on a low dimensional manifold in the space of all possible text data. If compression is optimised for reconstructing samples from a distribution with such manifold structure, then for observations that lie near the manifold the reconstruction will be drawn towards the manifold to an extent determined by available capacity. Furthermore, these generative models often contain an inductive bias, also characteristic of our text-VAE, that text is generated by independent latent factors of variation. Combined with natural text statistics this inductive bias results in the emergence of latent topics, which constitute clusters of semantically related words. Any given text excerpt is represented as a specific mixture of possible topics. When such an algorithm is used to encode word lists from the DRM paradigm, the underlying implicit assumption of the model is that the list was generated by some mixture of a few latent topics. As capacity is decreased, the reconstructed observation will become an increasingly prototypical exemplar of the activated topics, leading to the intrusion of semantically related lure words. We used the trained model to test this hypothesis on the DRM paradigm. For this, word lists of the original paradigm were taken as the set of words coming from a document and we inferred a posterior representation associated with this document. This posterior was subsequently sampled and the synthesised word list was taken as the reconstruction of the original word list (Fig 2A). As expected, since semantically related words are likely to occur together in natural text, the model improved recall accuracy of such word lists relative to lists of randomly selected words, however the price is the intrusion of non-studied but semantically related words into the reconstructed list (Fig 2A). The intrusions indicate that while there is a loss of information, the encoding and decoding process keeps the reconstructed observation consistent with a stored gist level interpretation of the original observation. Importantly, the frequency of the recall of lure words is similar to the average frequency of the recall of studied words, reminiscent of human performance in the DRM task (Fig 2B).

**Fig 2.**
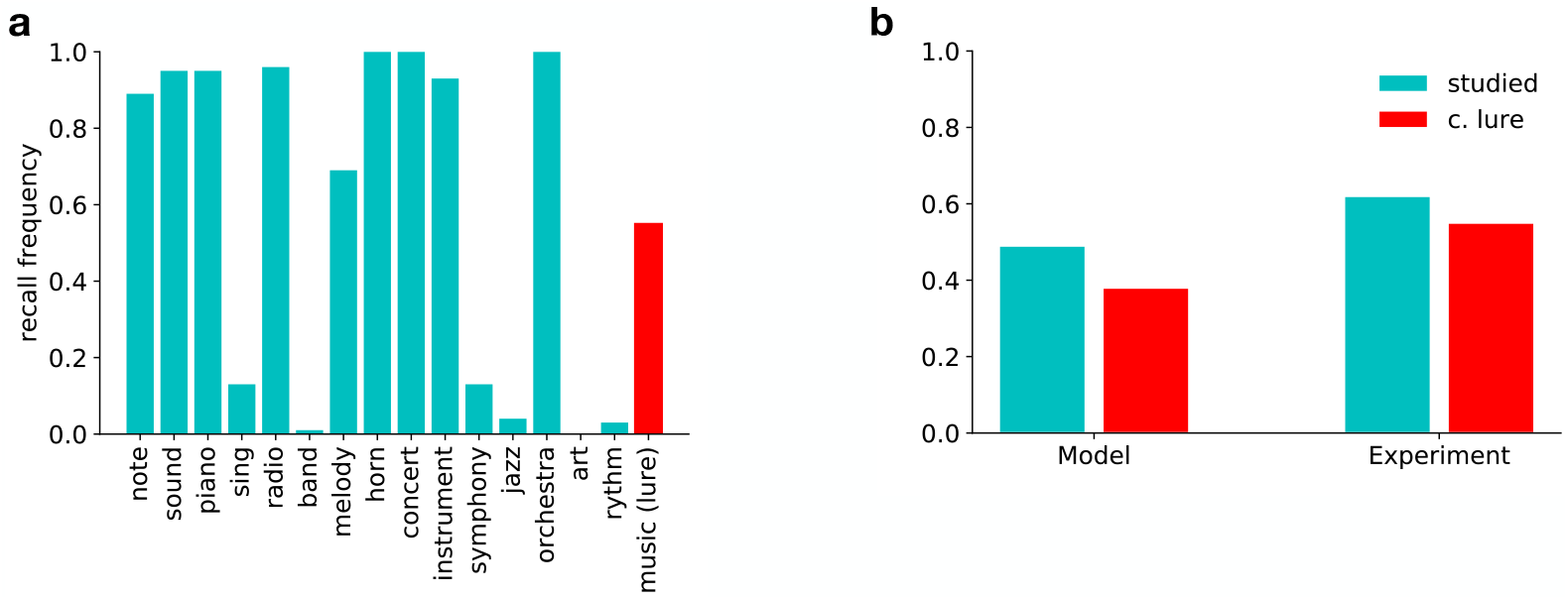
Memory distortions for lists of words. A: Frequency of recall from 100 samples for individual words in the text model trained on Wikipedia for the word list associated with ‘music’ from the DRM paper [39]. The lure word (*red*) is characterised by a recall probability comparable to the studied words (*green*). B: Comparison of recall probabilities for studied and lure words in the text-VAE model for 10 word lists (*left*, see Methods), and experiment (*right*) Roediger et al. [39].

#### Effect of varying contextual information on recall

We have argued that a second consequence of using the latent variables of a generative model for compression is that context influences both the degree and structure of distortions in recall. Hand drawn sketches of common objects have been used as complex naturalistic stimuli for exploring memory distortions, allowing the incorporation of contextual information by providing verbal labels or textual descriptions. The continuous nature of sketches allows us to explore graded and structured distortions of the observation along with the context dependence of encoding and recall. A dataset of millions of labelled sketch drawings created by a large and diverse set of human users of a browser-based game became available recently, and VAEs capable of handling these high dimensional data have been developed [41].

A well-known and robust example of the effect of contextual information is the experiment of Carmichael et al. [6]. In the classical experiment, intentionally ambiguous hand drawn sketches of objects from common categories were presented to subjects who were asked to reproduce these images after a delay. Two separate groups of participants were required to reproduce the sketches, with each group in one of two contexts. Context was established by providing a category name preceding the presentation of the drawings, with each name being consistent with one possible interpretation of the drawing. The authors found that, depending on the contextual cues, systematic biases were introduced in reproduced images that made the drawing more consistent with the provided label (Fig. 3B).

**Fig 3.**
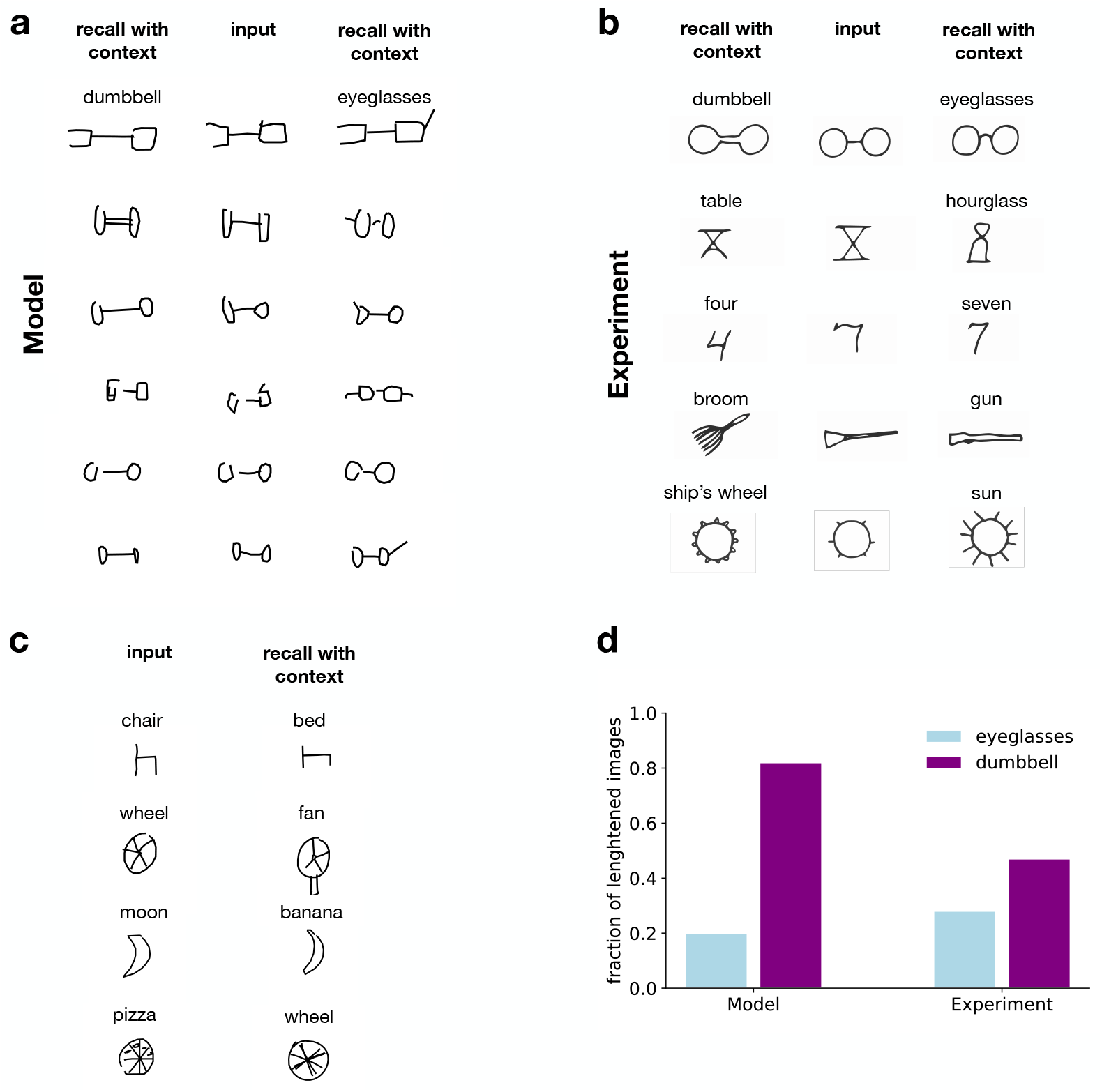
Context effects on reconstruction of line drawing from memory. A: *Middle column*, Ambiguous line drawings from the QuickDraw data set of eyeglasses and dumbbells. *Left* and *right columns*, Reconstructions of the image from memory in the dumbbell and eyeglasses contexts, respectively. Context is modelled by using a sketch-VAE trained on sketches from a single category with *beta* = 2. B: Examples ambiguous drawings (*middle column*) and their reconstructions (*side columns*) when cues are provided to participants (*shown as text labels*). Data is reproduced from [6]. C: Effect of contextual information on the visual features in recalled stimuli in the model. Quantitative changes (*top*), qualitative changes (*middle*), and subtle changes in characteristics (*bottom*) occur as a result of contextual recall. D: Quantitative changes in visual features with changing context (proportion of the length of the line connecting circular features in the eyeglasses and dumbbell contexts) in the Sketch-RNN model (*left*) and experiment (*right*). Experimental data reproduced from [42]

In order to analyse contextual effects in reconstruction in semantic compression we trained a VAE on sketch drawings from the QuickDraw data set (sketch-VAE). As an approximation of the semantic model for sketch drawings, we used the Sketch-RNN architecture, which models sketches not as raster images but as a series of sequential pen movements [41]. The model uses recurrent neural networks to make predictions on each subsequent stroke conditioned on its hidden state and the previous one and assumes Gaussian motor noise. We have selected ambiguous object-pairs from the QuickDraw data set and trained the model on 75000 drawings of each category. In order to model the effect of presenting a contextual cue, we have trained a conditional model on sketches belonging to each label. Each of these models represents a label conditional generative distribution *p*(*x*|*z*, *y* = *label*_*i*_) and a label conditional approximate posterior *q*(*z*|*x, y* = *label*_*i*_). During inference, we use the corresponding distributions to reconstruct the same ambiguous image. Consistent with human data presented (Fig. 3B), reconstructions from the conditional posterior resulted in systematic distortions of the original image consistent with the provided label (Fig. 3A). Systematic distortions introduced by the model were rich, spanning addition or deletion of features, rescaling of features, or subtle but characteristic changes in the shapes of reconstructed drawings (Fig. 3C). Systematic rescaling of features has been observed in humans [42], which is qualitatively similar to the rescaling found in the sketch-VAE model (Fig. 3D).

### Rate distortion trade-off

The value of retaining information from a given episode is likely to vary with respect to a multitude of factors such as how surprising the episode is, its relevance for predicting the near future or its emotional valence. As a consequence, we propose that memory resources allocated to storing episodes are unlikely to be constant either at the time of encoding or as a function of time. If memory resources are to be distributed rationally, this memory decay should not result in random forgetting as information theory provides a principled way of discarding information so that memories degrade gracefully.

Formally, optimal forgetting entails moving along the line of optimal encodings in the rate distortion plane in the direction of decreasing rate (Fig. 4A). At one extreme, where the rate distortion function intercepts the rate axis, resources are sufficient for lossless compression, meaning that verbatim recall is possible. At the other intercept, no information is retained relating to the individual episode and reconstruction is based purely on knowledge of environmental statistics. Starting from the point corresponding to verbatim compression, the memory trace becomes increasingly gist-like, until a point where even a very high level gist of the episode is lost. This way, the trade-off between rate and distortion results in the emergence of a continuum between gist and verbatim representations.

**Fig 4.**
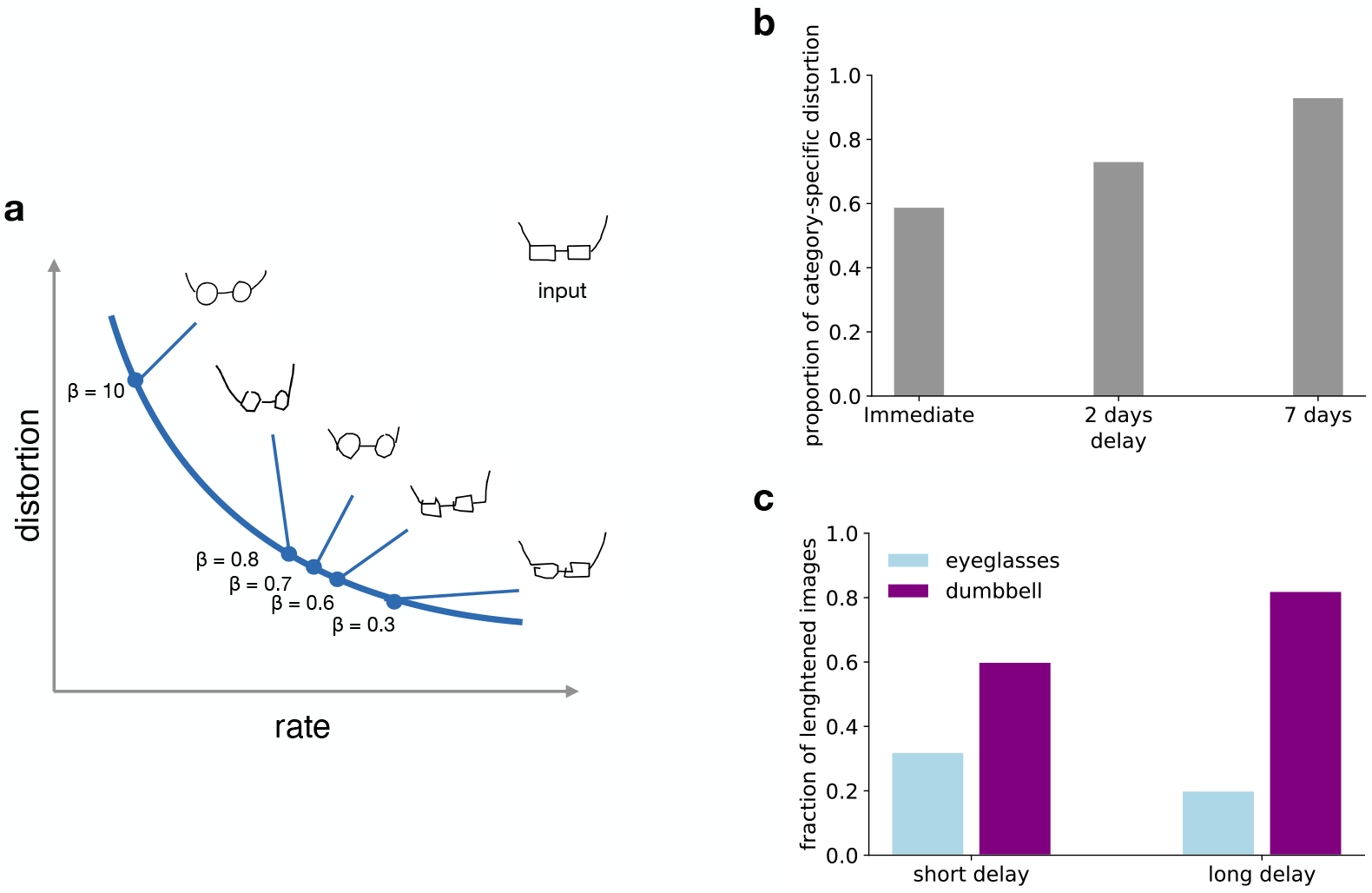
Rate distortion trade-off in memory for sketch drawings. A: Illustration of stimulus reconstructions as changes in *β* result in different points on the rate distortion curve for the sketch-VAE model. Inset image is used as input and is reconstructed with various levels of compression. Optimal forgetting implies moving along the curve in the direction of increasing *β* corresponding to increasingly prototypical reconstructions of the original drawing. B: Proportion of recalled sketches judged to show category specific distortions in humans due to the context presented during learning at different delays between stimulus presentation and recall. Distortions were evaluated by two of the experimenters and one judge naive to the purpose of the experiment. Figure reproduced from [42]. C: Proportion of sketches reconstructed by the model showing category specific distortion as a function of increasing compression. Quantitative changes in visual features are assessed, similar to Fig. 3D.

#### Temporal reduction of memory resources

In cognitive processes, such as prediction, time delay between storage of information and its retrieval is a factor that fundamentally affects its relevance and therefore the resources that should be dedicated to the particular piece of information. Anderson et al. [12, 43] proposed that it is a property of the natural environment that there is a decreasing need for information contained in individual traces as time progresses. For example, information contained in an email is more likely to be needed within a day of receiving a message than after a month. By studying library borrowings, access times of digital files, email sources and word appearances in the headlines of newspaper articles they concluded that forgetting curves demonstrate that human memory is adapted to this decreasing demand. These results have been corroborated by forgetting curves of US presidents in multiple generations of college students [44]. This argument motivates the idea that different rate-distortion trade-offs can be studied by controlling the time between stimulus presentation and recall or recognition. The effect of retention interval has been studied both in the recall of hand-drawn sketches and even more extensively in the DRM literature, allowing us to contrast it with our model’s predictions on targeting various points of the rate distortion trade-off.

In order to model the effect of delay, we have optimised models for increasing levels of compression by training them with increasingly larger *β*s. Since stronger compression implies more gist-like reconstructions, the high level context has a stronger effect on the recalled drawing as memory resources are decreased in the sketch-VAE model (Fig. 4A). We assessed the scaling of features for the ambiguous eyeglasses-dumbbell stimulus in the two contexts as a function of available memory resources. The analysis demonstrated more frequent category-related distortions for *β*s associated with longer delays. (Fig. 4C). Similarly, higher levels of context related distortions were found for the same pair when recollection was tested with increasing amounts of delay with human participants [42] (Fig. 4B).

Several studies have examined the effect of delay on recall performance in the DRM paradigm [45–47]. Toglia et al. [45] has performed the experiment with recall immediately after presentation of the lists or after delays of one or three weeks. In contrast with most of the prior work on the subject, retention intervals were varied between subjects, avoiding artefacts due to retesting the same subject. They have found that while recall for studied words had fallen sharply, recall of lure words was relatively unaffected even after three weeks. In a variation of the original paradigm, for some of the subjects they have presented lists in a ‘random’ condition, pooling the words from six lists and presenting them in shuffled order. Interestingly, in the random condition recall probability of lure words had increased as compared to shorter delays at week 3. An even longer delay of two months has been studied by Seamon et al. [46], where they have found that eventually the recall of lure words also approaches zero. Thapar & McDermott [47] have looked at a similar design with a maximum delay of one week while also modulating depth of processing at the time of encoding. Many other studies besides these have examined the effect of various manipulations on accuracy for delayed recall but it has been a robust finding that memory for lure words, that is false memories, are more persistent in time than memory for studied words (Fig 5D).

**Fig 5.**
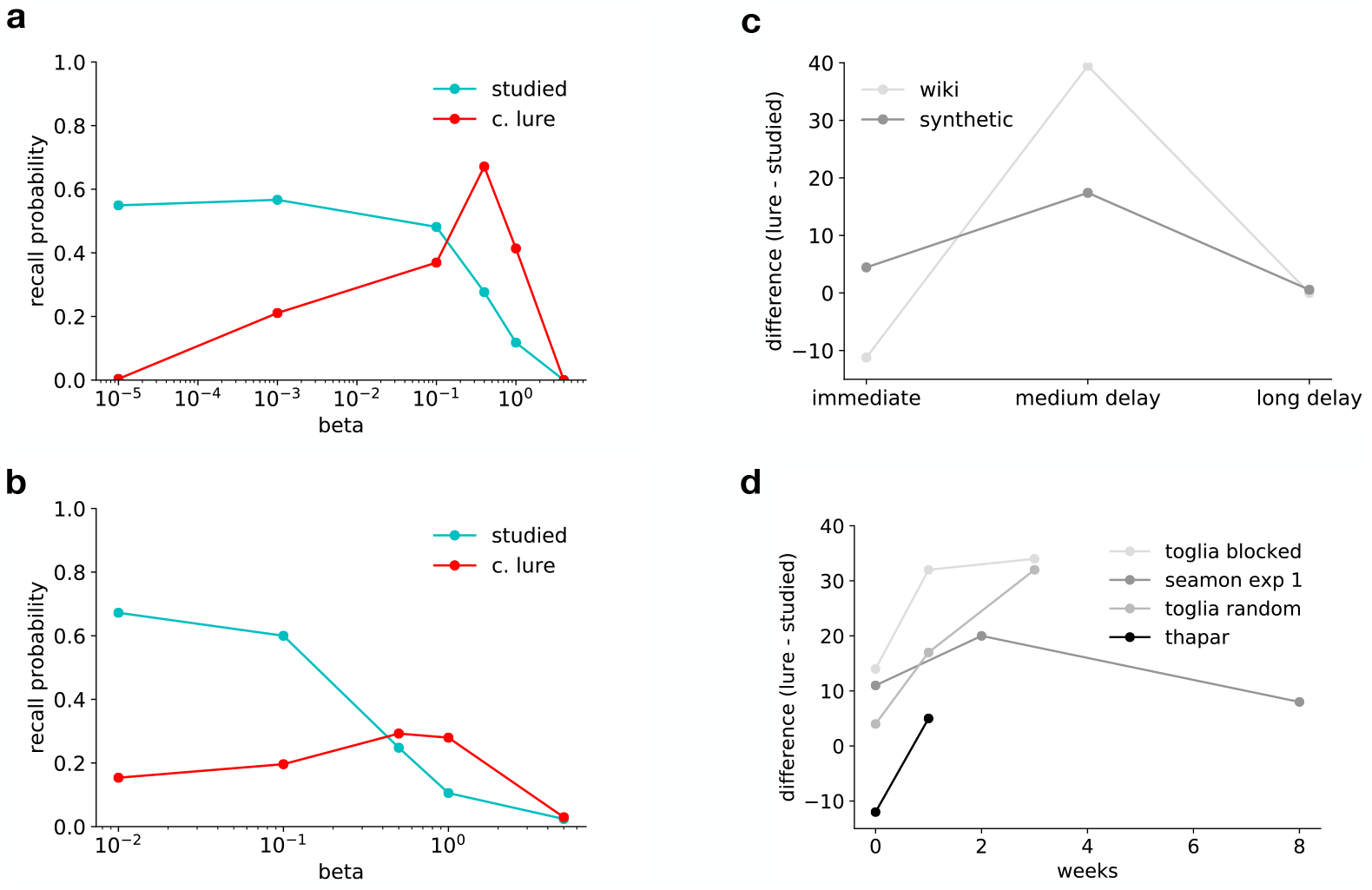
Rate distortion trade-off in forgetting over time. A,B: Using the text-VAE model, we modelled the dependence of memory representations on time by gradually increasing the compression rate, *β*. Recall probabilities averaged over multiple word lists of studied and lure words was measured on a text-VAE trained on Wikipedia entries (*top*) and on a synthetic vocabulary (*bottom*). Increasing the compression rate results in a monotonically decreasing recall performance for studied words. In contrast, increased delay of recall leads to an increase in false memories. For critical NS (lure) words the recall probability initially increases with larger compression rates but very high compression rates result in losing gist-like recall as well. Asymptotically the performance on semantically related S words will approach the performance on random word lists as less and less of the structure of the data is used. C,D: Difference between recall probabilities for lure words and studied words as a function of the delay between recall and study for the model (*top*) and experiments (*bottom*). Both Wikipedia-trained and synthetic vocabulary trained models predict persistence of false recall of non-studied lure words as compared to studied words, visible as an increase in the difference between lure and studied word recall rates as a function of time. For even longer delays, the gist information is progressively forgotten as well and consequently recall rates for both lure and studied words approach zero. The same pattern of increasing rate up to a delay of three weeks and a subsequent decrease can be observed in experimental data. Data is reproduced from Toglia et al. [45], Seamon et al. [46] and Thapar & McDermott [47].

We have investigated the dependence of recall on memory constraints in the word list recall paradigm using our text-VAE model. We have trained separate models for each memory constraint, corresponding to various levels of the rate distortion trade-off parameter *β*. We then used these models to reconstruct each word list at each setting of the memory constraint and subsequently evaluate recall accuracy for studied and lure words separately. Reflecting a transition from a verbatim-like to a more gist-like latent representation, recall performance on studied words decreased monotonically with increased level of compression. The decrease in accurate recall was paralleled by an increase in gist-based intrusions of related lure words starting from a negligible level as the latent representation incrementally came to resemble a topic model. This was followed by a reversion of false recall rates to negligible levels, reflecting a loss of even gist information due to extremely limited capacity (Fig. 5A). Note that we have restricted our analysis to the main effect of length of delay in the delayed recall DRM experiments and thus our text-VAE does not explain the effects of manipulating depth of processing or word order effects. In order to control for the mismatch between Wikipedia and natural text statistics, in addition to the text-VAE trained on Wikipedia articles, we also tested model performance on synthetic data. We generated a synthetic corpus with controlled statistics from a Latent Dirichlet Allocation (LDA) topic model, which models statistical structure in text by assuming the presence of latent topics that are present in each document. In order to construct DRM lists for the synthetic model, we have used nearest neighbours according to word embeddings learned from the synthetic corpus (see Methods for further details). The synthetic data trained text-VAE shows a transient increase in lure recall, similar to the one observed with the Wikipedia-trained model (Fig. 5B).

We have argued that a decreasing need for information contained in a memory trace over time implies that memory resources assigned to the trace should be decreased. Therefore, according to RDT, the delay between encoding and recall corresponds to a change in the RD trade-off, controlled by the *β* parameter. However, the exact mapping between length of delay and the numerical value of *β* can’t be derived from the theory but has to be calibrated based on measurements. In the standard DRM experiments, recall probabilities of studied and critical non-studied words are similar (Fig 2D), and consequently we set *β* corresponding to immediate recall to be the same as in our model of the original experiment (*β* = 0.1 for the Wikipedia trained model and *β* = 0.5 for the model trained on synthetic data). Increasing the rate from this level (decreasing *β*) results in increasing accuracy and thus the gradual disappearance of false memories. Since time delays are necessarily positive, the predictions of the model in this regime describe a hypothetical situation where memory resources available to the subjects were increased above what is typically measured in this paradigm. On the other hand, modelling memory decay in time by decreasing the rate results in a decrease in the recall of studied words, and an initial increase (for the models trained on Wikipedia) or much less pronounced decrease (for synthetic data) in false memories, resulting in a similar pattern of relative advantage of lure words over studied words after medium-length delays as the one seen in human experiments (Fig 5C). At very high levels of compression (high *β*), recall for all word types is poor, as even the broad theme of the list becomes forgotten and the model samples lists of related words that occur together frequently in natural text.

## Discussion

In this paper we demonstrated that the principle of lossy compression provides the basis for a unifying normative account of a wide variety of systematic memory errors. Central to our approach was that we related inference in probabilistic generative models to rate distortion theory: inferring the latent variables underlying observations amounted to selectively discarding information not represented in the latent variables, thus establishing a lossy compression method. We used a recently developed machine learning framework to train a probabilistic generative model on a variety of naturalistic, high dimensional data sets (chess games, line drawings, and natural text). We showed that the effect of domain expertise on recall accuracy in remembering chess board positions and gist-based distortions in remembering semantically related word lists arise as straightforward consequences of optimising the rate distortion objective. Furthermore, we demonstrated the emergence of varying degrees of ‘gistness’, resulting from the rate distortion trade-off.

### Interpreting memory distortions as lossy compression

The experiments discussed in the study demonstrate key consequences of semantic compression in a single computational framework. The primary appeal of this framework is that it can integrate a large variety of experimental observations under a simple computational principle. The demonstrated effects, however, are more general than the experiments discussed here. For instance, the effect of domain expertise has widespread support in the memory literature not just in the domain of chess but also memory for sports trivia, software code, medical images, and other games [48]. In addition to the effect of varying levels of expertise on recall accuracy, a similar effect arises if the congruence of stimuli to the statistical structure of the domain is varied along a spectrum, for example the order to which letter statistics of words conform to that of the English language [49]. The encoding of the observation into a posterior over latent variables can be understood as compressing sensory experience into sufficient statistics for the latents, which in addition to the gist-based distortion experiments analysed here explains seemingly paradoxical results that discrimination performance of sound textures decreases with increasing stimulus duration when stimulus samples come from the same texture family [50]. Such a process also implies that the level of difficulty of inference affects the accuracy of the recalled memory trace. Similar to the Carmichael effect, classical memory experiments have shown that providing a concise context which aids the interpretation of otherwise strongly ambiguous stimuli can greatly increase retention accuracy [5, 7]. Beyond the effect of expertise on reconstruction accuracy, the Chase and Simon experiments (1973) also display effects that are related to representing latent variables. In particular, temporal dependencies in placements of chess pieces were suggested to reflect chunking mechanisms. The proposed variational autoencoder framework naturally generalises to these domains as well.

The delayed DRM paradigm has been extensively studied [45–47]. In this study we only have addressed the effect of delay alone but effects such as divided attention, order effects and depth of processing were not discussed. Some of these could be addressed by natural extensions of this model, for example order effects would require extending the model to non-iid observations. Others, such as the effect of varying depth of processing during encoding or recall we do not see as direct corollaries of the RD framework and therefore would require further assumptions.

We have modelled forgetting as the effect of decreasing capacity allocated to a memory trace over time by training separate models for different levels of compression. A limitation of this approach is that there is no guarantee that a slightly compressed representation taken from a model with low *β* can be converted into the strongly compressed representation of the model given by a high *β*, as it is possible that the high *β* representation utilises information that is present in the original observation but not in the low *β* representation (although in toy settings this seems not to be the case as the introduction of further capacity leads to capturing additional data generative factors [51]). Furthermore, taken as a process model this approach would require storing a separate semantic model for each available level of compression. Approaches where a single model is trained for compression at multiple rates have been proposed in the machine learning literature [52, 53]. In addition, hierarchical generative models have been introduced that are capable of generating observations at progressively increasing level of detail by conditioning on variables at more and more levels of the hierarchy [54–56]. Such hierarchical generative models define a straightforward process for converting a memory trace into one requiring lower capacity by selectively discarding information. The lowest level of compression is achieved by storing all levels of latent variables, then as time progresses, the states of successive levels of variables are discarded beginning with the one closest to the input layer. Some of these models show semantically meaningful partition of information between the layers in limited domains. For example in Maaloe et al. (2019), the compression process outlined above applied to portraits initially retains information about wearing glasses but discarding the specific information that those are sunglasses, and at later stages of compression it forgets about the glasses while keeping a large portion of facial features still intact. One way in which such a compression could be implemented is if semantic compression utilises the hierarchical representations in sensory cortices as have earlier been argued in [57–59].

### Theoretical considerations

Application of RDT to lossy compression in the brain seems to be a natural choice. A closer inspection of the problem, however, reveals that from the perspective of compression and specifically its formal theory, RDT, a fundamental challenge arises. In memory systems, the data set used to learn the model is inherently incomplete, that is only a subset of the data that specifies the model had been observed. This setting defies a critical assumption of RDT that the data statistics are known. The brain tackles the issue of incomplete data by updating the model continuously when new data is observed. However, the constraint that the statistics of observations is being learned concurrently with using it for compression places unique demands on a memory system. We have previously argued that if the overall structure of the model describing the environment is known then the parameters of the model can be updated once new observations are made and the only information that needs to be maintained is the sufficient statistics of model parameters. Consequently, other features that are not part of the sufficient statistics can be discarded without harm. However, if there is uncertainty over model structure and it is not *a priori* known which features are relevant for the model, then the ability to reconstruct the data becomes critical [31]. The need for such reconstructive ability motivates our use of unsupervised learning, which attempts to capture all of the variance in the data when resources are not bounded, constituting a perfect episodic memory. In case that relevance has to be sensitive to predictive ability [60, 61], rewards or a supervision signal, the framework can be straightforwardly extended through the same deep variational information bottleneck objective [32, 62]. Task variables can potentially also be accommodated in a generative perspective in which task variables are part of a generative model. Such models have been introduced in machine learning [63].

The variational approximation that we have used here provides a useful tool for integrating principles of RDT and probabilistic generative models, which can be tested under conditions where data complexity is close to that of the natural environment. Since this is the data set that human memory systems are adapted for, we believe that these are relevant stimuli to contrast capacity constraints of the model and that of human memory systems. Application of the framework, however, also comes with specific choices and alternative formulations are possible. In RDT and the IB method, the rate term is defined as the mutual information between the observation and the latent code which the variational method provides an upper bound on. This is an abstract constraint, and the specific influences on the latent representation depend on model architecture. For example changing the Gaussian prior to a Laplace distribution would result in a constraint on the sparsity of latent activations. Furthermore, it has been argued that memory resource constraints for the brain would be better captured by restricting the representational cost of storing the encoding, corresponding to minimising the entropy of the code instead of the mutual information [64]. This choice leads to the Deterministic Information Bottleneck (DIB) method, which can lead to qualitative differences such as the optimal reconstructions being deterministic.

Variational approximations also exist for the DIB, and it has been argued that a discrete latent space variant of variational autoencoders called Vector Quantised Variational Autoencoder can be viewed as an approximation to the variational DIB principle. Furthermore, the correspondence between RDT and generative models can be drawn in alternative ways: Balle et al. [34] show a correspondence where the posterior is of fixed variance and the multiplication factor beta arises from the variance of the observation noise. We see the information theoretical form of the bottleneck constraint as an approximation to multiple constraint terms, possibly arising from the demands of cognitive functions other than memory, each having a contribution to shaping the latent representation. The question of which variant of the computational framework and what combination of constraints would correspond most closely to representations and memory distortions measured in human experiments is a subject for further investigation.

### Related work

RDT has recently been proposed as a framework to investigate distortions of memory by Bates and Jacobs (2020) [65]. While the computational principles they apply and those in this and our previous work [66] have strong parallels, the differences highlight different aspects of using generative models for compressing complex data. While VAEs constitute state-of-the-art in machine learning for learning generative models of high dimensional data, even these models struggle to capture the full richness of natural stimuli. In particular, learning highly structured noise models has proven difficult, leading to issues such as blurred reconstructions [67]. In order to mitigate this problem, instead of using pixel image data, we have opted to use data represented in low level features such as chess board locations or pen stroke endpoints. This choice essentially circumvents the problem of learning the low level noise model for the network. Bates and Jacobs [65] take the alternative approach of working directly with pixel data and thus provide an end-to-end learned model. The choice of training the model end-to-end versus learning over low level abstract features has complementary benefits: while using pixel data to study memory effects is certainly appealing since the perceptual process is more completely integrated, it has the disadvantage that the generative model needs to cope with limitations of VAEs on natural images, such as blurry reconstructions. Blau et al [68] proposed that the issue of perceptual quality of reconstructions could be mitigated by introducing an additional term to the RD trade-off, which could be optimised utilising the framework of the other prominent form of deep generative models besides VAEs, Generative Adversarial Networks [69].

We have argued that while we rely on the unsupervised version of the information bottleneck to make the connection between RDT and latent variable models, the information bottleneck is originally framed as a supervised method targeting the relevant information regarding a task variable and the beta-VAE extends naturally to this setting through the same variational objective [32]. Consequently, RDT enables incorporating the effects of task demands on the learned representation in a principled way through the specification of the distortion function. While introducing such demands into the distortion would allow an exploration of further aspects of memory distortions, we left these for future work, restricting ourselves to the unsupervised version in our study. Bates and Jacobs propose another method for incorporating task-variables in their model by extending the unsupervised component with a decision network, allowing task-variables to affect the latent representation. Among other applications, they use these task variables to model category bias, however they point out that such bias can also appear due solely to categorical structure present in the data distribution. This latter explanation is what we appeal to in our study, although we agree that the latent representations in semantic memory are presumably also shaped by task objectives. We believe that further progress in machine learning in the area of learning generative models will allow lifting current limitations and will provide the background for a fully consistent model of memory.

Compression, and more specifically RDT has been proposed as a framework for an ideal observer analysis in visual working memory and perception tasks [70–72] In Sims et al. [70] they experimentally demonstrate RDT’s prediction that if memory is optimised for the statistics of stimuli learned in the course of the experiment, recall should be less accurate in case the distribution of stimuli has high variance as opposed to a low variance condition. In Sims et al. [71] they infer the distortion function in a bottom-up fashion from behavioural data. One major point of contrast between this approach and ours is that instead of inferring the distortion function from behaviour, in our study the distortion function is implicitly defined by the inductive biases inherent in using latent variable generative model used for compression. A second point of contrast is that in all of these works, the authors apply RDT to low dimensional perceptual tasks with simple statistics, where optimal encodings are feasible to compute directly. In our approach it is the complex, high-dimensional and strongly structured nature of input statistics that necessitates the use of generative models which in turn define the distortion function. In [72] they analyse colour perception and memory and in addition to a low-level distortion term measured in pixel space they introduce an additional term which penalises distortions that result in the reconstruction crossing colour category boundaries. The additional term in the distortion introduces a category dependence of reconstruction similarly to the category related distortions we have modelled with the sketch-VAE, however they did not explore contextual effects in reconstruction. We argue that these perceptually simple tasks, while allowing for quantitative comparison of predictions with experimental data, are less capable of inducing the kinds of distortions that we are concerned with here, as the need for generative models is most crucial when the input distribution is complex, high-dimensional and strongly structured.

Our work is closely related to Bayesian account of reconstructive memory approach of Hemmer et al. [17] where they provide a normative method for combining episodic and gist information available in memory through an optimal Bayesian decoder. They assume that stored values for features in the memory trace are noisy versions of the observed values. These noisy values are then combined with feature priors through Bayes’ rule, which reduces noise in the memory through exploiting prior knowledge. This decoding step is similar to reconstructing the observation in a generative model conditioned on inferred latents in our approach, however they do not consider the encoding step as part of the same process which should be optimised. In semantic compression, features are prioritised according to the distortion function and the optimisation also concerns what information should be kept as part of the trace. As a result, the amount of memory noise can vary as a function of how important each feature is in relation to others and the memory constraints, which is a fundamental difference from the setting considered in Hemmer et al. [17] (e.g. the sketch-VAE automatically chooses a trade-off in accuracy between features such as the presented glasses’ shape, the angle of the rims and the length of the bridge connecting the rims). Hemmer & Steyvers [30] use a dual-route generative model to explain the effect of semantic memory in a scene recall task, but they do not relate their method to compression. Their topic model of semantic memory is very similar to our text-VAE for certain settings of beta, and they have a parameter corresponding to capacity. However, this capacity parameter only affects the episodic route and does not affect the representation of the semantic model. A crucial contribution of the RD perspective is that it provides a principled way of changing the representation as a function of available capacity. As a result, in our treatment dual routes are not required, since RDT provides a continuous trade-off between episodic-like and semantic-like memory traces. Furthermore, as the reconstructive ability of the semantic route is not affected by capacity in their model, we believe that explaining delayed recall results of very long delays such as in Seamon et al. [46] would require even further assumptions. A trade-off similar to that implied by RDT but without establishing a formal link to the theory of lossy compression has been formulated in the context of communicative interaction and leads to the emergence of semantic categories [73].

Human memory has a remarkable capacity to adaptively support decisions in a versatile environment but it also displays a rich array of distortions [13, 74]. These systematic errors have the potential to shed light on the design principles of our memory systems [12]. Performing complex tasks by agents suggests that various computations can be supported by episodic and semantic memory systems [75–78]. Accordingly, memory distortions have also been linked to different computational processes. In particular, besides diminishing resources, other normative arguments have been made to understand various aspects of time-dependent deterioration of memories. Interference of memories from novel experiences has been linked to the flexibility of the represented model [79, 80] and regularisation was proposed as a normative principle, which could help preventing overfitting [81]. Dynamics of the environment has been linked to adaptive forgetting rates [82, 83] and destabilisation of earlier memories after their reactivation has been linked to model update [84]. Normative models of memory distortions fall into two broad categories. One family of studies explored how the usage of latent variables to encode experiences, i.e. performing inference in the internal model, introduces systematic distortions [30, 85, 86]. Another family of studies explored how updating this internal model of the environment, i.e. learning the internal model, leads to various forms of memory distortions [31, 87, 88]. Common in all these models is that an internal model of the environment is assumed to underlie learning and inference and these internal models are described in terms of a generative model of the environment. This indicates that capitalising on more complex generative models capable of learning representations of more naturalistic data in multiple domains can contribute to a deeper understanding of memory dynamics in natural environments.

## Materials and Methods

### Chess-VAE

We have trained a beta-VAE on games downloaded from the FICS (Free Internet Chess Server) Games Database, containing hundreds of thousands of recorded online games. Chess configurations were represented as a one-hot vector for each one of 64 squares on the chessboard, with each 13 dimensional one-hot vector specifying the chess piece or the lack of a chess piece in that square. The decoder output was a categorical distribution for each square, which represented the probabilities of possible chess pieces on the particular position. Both the encoder and decoder of the chess-VAE consisted of two dense layers with 3000 units and sigmoid nonlinearity, followed by a linear transformation. The prior over the continuous 64 dimensional latent space was a Normal distribution with an identity covariance matrix. Reconstruction was modelled by conditioning the model on a chess configuration, inferring the latent representation, then taking the MAP reconstruction of the decoder.

Since the goal was to demonstrate robust qualitative effects resulting from the theoretical framework, hyperparameters of the model were chosen so that reconstructions would fall in a regime comparable to the experimental data as opposed to fine tuning them for an accurate match. Beta is a central free hyperparameter for the model, which was chosen to be 0.0001 so that the ‘expert’ model could reproduce the state of around 90% of squares correctly in the MAP estimate. Decreasing beta further did not result in flawless recall, presumably due to capacity limitations in the encoding and decoding transformations. Patterns in the presented results were relatively robust to changes in parameter *β* but increasing it by orders of magnitude causes both the overall accuracy and difference between the random and game conditions to decrease. Varying the size of latent space had similar effect to varying *β* as it modulates the capacity of the latent space. Decreasing the hidden layer widths from 3000led to qualitatively similar results but with performance in the game board condition saturating at a lower level of around 15 successfully reconstructed pieces in the case of 1000 units and 10 in the case of 500 units. Doubling the number of training steps did not noticeably change the resulting reconstruction accuracies. . Separate training and test sets were formed from all the board positions from 3000 games of the FICS Games Database. Different levels of skill were modelled by using different sized training sets: game positions were subsampled to 0.1%, 1%, 10%, and 90% of the full dataset. Training set sizes corresponding to various skill levels were also not precisely calibrated to experimental data but set to what we determined to be sensible values so that amateur players would only observe a few games and expert players would see a sufficient amount to reconstruct with comparable accuracy to the experiments. Remaining set sizes were chosen to span orders of magnitude between these two extremes. For training we have used Adam with a learning rate of 10^−4^ and a batch size of 65.

The test configurations were constructed as in Chase et al. [89]. ‘Game’ configurations were taken after either the 41st move in games from a separate test set. ‘Random’ configurations were produced by shuffling the pieces across occupied board positions of a game setting. The number of pieces on the board were not fixed in the dataset and could vary across trials. We have followed the accuracy evaluation method proposed in the Gobet and Simon paper, where the number of correct pieces on a reconstructed board is counted.

### Text-VAE

The beta-VAE constructed to learn the statistics of natural text was similar to the chess-VAE: we used an encoder and a decoder with two dense layers with 2000 hidden units per layer followed by sigmoid nonlinearities. Activations, **z**, in the last hidden layer of the decoder are used to generate words, **X** independently through a linear transformation and a softmax nonlinearity according to

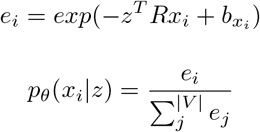

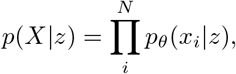

where R is a |*z*| × |*V*| matrix that acts as a semantic word embedding, organising words into a low dimensional continuous vector space such that similarity in this space reflects semantic similarity between the words. The model had a continuous latent space with a diagonal Gaussian prior in 100 dimensions. This architecture (with minor differences) has appeared in the machine learning literature previously as the Neural Variational Document Model (NVDM) [40]. The factorisation assumption, which formulates the generation of multiple-word text as an independent process, greatly simplifies training but the consequence is that the same word can be generated multiple times in a synthesised text. Since this is a purely computational simplification that is clearly detrimental to performance in the world list recall task, we constrained reconstructions such that distinct words had to be generated. For training on both Wikipedia and the synthetic dataset described below, we have used Adam with a learning rate of 10^−5^ and a batch size of 100. We have chosen these hyperparameters based on the Miao et al. [40] paper, adjusting them for the fact that they have trained their model on significantly smaller datasets (vocabulary sizes of around 2000 and 5000 as opposed to 50000). The large dataset was required so that we have a sufficiently good approximation of language statistics, which we assessed through testing whether the associations that the original DRM lists rely on are reliably represented in the model. Consequently we have increased the training set to the extent that was computationally feasible for us to train. Beyond these considerations no further adjustment of the model parameters was required and results are presented without fine tuning of the parameters. Due to the computational cost of training the model, we have not explored perturbations of hyperparameters extensively. We have calibrated *β* values such that the reconstruction probabilities for studied and critical lure words were approximately the same, which is what was observed in the original DRM experiment.

To control for the mismatch between the statistics of the training corpus and the natural vocabulary humans experience over a lifetime, we only used the word lists from the original DRM article which fulfilled the criterion that the lure word had at least 2000 occurrences in the training set. Words that appeared less than 100 times in the training set were deleted from the lists, and if these manipulations resulted in a list that was shorter than 12 words then the entire list was removed. According to these criteria we included 10 of the original word lists in the analysis (high, rough, mountain, music, black, man, foot, king, river, soft). As an illustration of the similarity of associations between the model and human data, 15 closest associates of the word ‘music’ in the model are ‘musical, album, arts, songs, sounds, pop, art, instrument, sound, musicians, progressive, disambiguation, label, composers, string’ whereas the corresponding DRM list is: ‘jazz, horn, concert, orchestra, rhythm, sing, piano, band, note, instrument, art, sound, symphony, radio, melody’.

In order to mitigate variability in the averages due to i) the low number of word lists and ii) mismatch between the statistics of the Wikipedia training set and natural text, we also built a synthetic dataset so that the performance of the text-VAE can be explored under well controlled statistics. We generated synthetic text using an artificial vocabulary and a Latent Dirichlet Allocation (LDA) topic model. The LDA generative model used a synthetic vocabulary of 1000 words with a word concentration parameter 0.1, and 10 topics with concentration parameter 0.1. These parameters were chosen such that a t-SNE embedding would largely be able to separate the main topics in each document but not perfectly. We sampled 20000 documents from the LDA models to use as a training set. In the original memory experiment, word lists were generated by asking subjects to list their first associates to the presented lure word. Analogously, for the artificial vocabulary we trained a separate model on synthetic data generated from the LDA model and computed the 15 most similar associates based on the learned embedding matrix R. We generated such DRM-like word lists for each word in the vocabulary that occurred at least 500 times in the training set, but discarded lists that became shorter than 12 after removing infrequent (less than 500 occurrences) words. Note that this method can also be used to construct new DRM word lists based on the model trained on Wikipedia.

### Sketch-VAE

To obtain a generative model for hand-drawn sketches we used the sketch-RNN VAE architecture [41] that was developed specifically as a generative model for sketches and is able to capture sequential dependencies in the data. Sketch-RNN represents sketches as sequences of pen strokes, which are encoded into the latent representation through a bidirectional RNN. The output of this network then parametrises a Normal distribution over the latent space. The decoder consists of an RNN conditioned on the latent vector and preceding strokes, outputting the parameters of a mixture of Gaussians which generates the next pen movement.

In order to be able to relate our analysis to the distortions observed in the Carmichael experiment, we have used hand drawn sketches from the QuickDraw dataset Ha & Eck [41], consisting of a rich set of labelled drawings depicting 345 common object categories. The experiment used ambiguous drawings that could plausibly belong to multiple categories; hence we selected category-pairs that contained a substantial number of visually similar exemplars. Although the QuickDraw dataset contains rich naturalistic samples from every category, characteristics of recording the doodles preclude a large number of object pairs from the analysis. The QuickDraw data was recorded as part of a web browser game, where subjects had 20 seconds to draw an exemplar of a given category. However, if the drawing gets to a stage where an object classifier is able to recognise it as belonging to the provided category, a new trial is initiated. As a result, the data set contains a large number of unfinished drawings. Another limitation of the data set is that participants tend to draw prototypical exemplars of the category thus limiting the variance of the samples compared to all possible ways of drawing the object that would still be easily recognisable by human observers. This means that some of the designs appearing in the Carmichael experiment are not present in the data set and thus the model is oblivious to their interpretation. In total we have selected five object pairs (eyeglass-dumbbell, chair-bed, wheel-fan, moon-banana, pizza-wheel) and the model was trained separately for each object category, on a training set of 75000 samples per category. The reconstructions were based on samples from a separate test set. We have modelled the effect of presenting the label by conditioning the category specific generative model on the ambiguous image and using it to generate a reconstruction.

Most parameters of the sketch-rnn model are determined by the data statistics and only a small subset of the parameters is available for fine tuning. These parameters have been explored in the original publication of the paper [41]. We have only explored the rate distortion parameter *β*. The value of *β* for different time delays were chosen so that the variance in reconstructed images was roughly comparable to that observed for humans in the experimental literature, as judged by the authors.

In the quantitative analysis of feature rescaling, following Hanawalt et al. [42], we have measured the proportion of the length of the drawing and the length of the connecting line for conditional reconstructions for 50 randomly selected samples of the test set. The length of the drawing was measured between the widest extent, excluding any stems in the case of glasses. The connecting line was defined as the distance between the two circular features at the points of intersection with the connecting line. We have only included samples where both the circular features and the connecting line was recognisable for all reconstructions.

## Acknowledgements

The authors would like to thank Mihály Bányai and Csenge Fráter for discussions and comments on the manuscript and to the anonymous reviewers for their constructive criticism. This work has been supported by the National Research, Development and Innovation Fund of Hungary (Grant No. K125343).

## Author contributions

D.G.N., B.T. and G.O conceived the experiments, D.G.N. designed the experiments, D.G.N. performed the experiments, D.G.N. analysed the data, D.G.N. and G.O. discussed results, D.G.N. and G.O. wrote the paper.

## Footnotes

1 https://www.ficsgames.org

